# Persistent homology demarcates a leaf morphospace

**DOI:** 10.1101/151712

**Authors:** Mao Li, Hong An, Ruthie Angelovici, Clement Bagaza, Albert Batushansky, Lynn Clark, Viktoriya Coneva, Michael Donoghue, Erika Edwards, Diego Fajardo, Hui Fang, Margaret Frank, Timothy Gallaher, Sarah Gebken, Theresa Hill, Shelley Jansky, Baljinder Kaur, Philip Klahs, Laura Klein, Vasu Kuraparthy, Jason Londo, Zoë Migicovsky, Allison Miller, Rebekah Mohn, Sean Myles, Wagner Otoni, J. Chris Pires, Edmond Riffer, Sam Schmerler, Elizabeth Spriggs, Christopher Topp, Allen Van Deynze, Kuang Zhang, Linglong Zhu, Braden M. Zink, Daniel H. Chitwood

## Abstract

Current morphometric methods that comprehensively measure shape cannot compare the disparate leaf shapes found in seed plants and are sensitive to processing artifacts. We explore the use of persistent homology, a topological method applied across the scales of a function, to overcome these limitations. The described method isolates subsets of shape features and measures the spatial relationship of neighboring pixel densities in a shape. We apply the method to the analysis of 182,707 leaves, both published and unpublished, representing 141 plant families collected from 75 sites throughout the world. By measuring leaves from throughout the seed plants using persistent homology, a defined morphospace comparing all leaves is demarcated. Clear differences in shape between major phylogenetic groups are detected and estimates of leaf shape diversity within plant families are made. This approach does not only predict plant family, but also the collection site, confirming phylogenetically invariant morphological features that characterize leaves from specific locations. The application of a persistent homology method to measure leaf shape allows for a unified morphometric framework to measure plant form, including shape and branching architectures.

## Introduction

As generally flattened structures, leaves provide a unique opportunity to quantify morphology as a two-dimensional shape. Local features (such as serrations and lobes) and general shape attributes (like length-to-width ratio) can be measured, but numerous methods also exist to measure leaf shape more globally and comprehensively. A popular method to quantify leaf shape is to place (*x*, *y*) coordinates, known as landmarks, on homologous features that are related by descent from a common ancestor on every sample (Bookstein, 1997). The set of landmarks from each leaf can be superimposed by translation, rotation, and scaling using a Generalized Procrustes Analysis (Gower, 1975). Once superimposed, the Procrustes-adjusted coordinates of each shape can be used directly for statistical analyses. Landmark analysis excels in its interpretability, because each landmark is an identifiable feature with biological meaning imparted by the shared homology between samples. Because landmarks are homologous features, their use often reveals genetic and developmental patterns in shape variation (Chitwood et al., 2016a).

Not all leaves have obvious homologous features that can be used as landmarks. Further, when comparing leaves with disparate morphologies (e.g., simple vs. compound leaves), there may not be identifiable homologous points. Nearly all leaves have homologous landmarks at the tip and base, but if there are no other identifiable landmarks, an equal number of equidistant points on each sample between the landmarks can be placed (Langlade et al., 2005). The denser and more numerous such pseudo-landmarks are, the closer they come to approximating the contour itself.

Another method, the use of Elliptical Fourier Descriptors (EFDs), measures shape as a continuous closed contour, and can also be used when homologous features are absent. EFD analysis begins with a lossless data compression method called chain-code, in which the direction to move from one pixel to the next is recorded as a chain of numbers (where each link in the chain *α* is an integer between 0 and 7 specifying the pixel direction (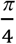) *a*) so that from this chain of numbers the closed contour can be faithfully reproduced (Freeman, 1974). The chain code is decomposed by a Fourier analysis into a harmonic series that is used to quantify an approximate reconstruction of the shape (Kuhl and Giardina, 1982).

Both pseudo-landmarks and EFDs measure leaf shapes for which homologous features that can be used as landmarks are lacking (Bensmihen et al., 2008; Chitwood and Otoni, 2017). Still, when comparing disparate leaf shapes, unless major sources of shape variance in the data (such as the number of lobes or leaflets) are present in every sample, individual pseudo-landmarks or harmonic coefficients will not correspond between samples in a comparable way useful for analysis. Recently, a computer vision method coupled with machine learning was used to classify leaves, with diverse vascular patterns and leaf shapes, into plant families and orders (Wilf et al., 2016). This method uses a visual descriptor to train a classifier. Since cleared leaves are used, this method relies on both internal features like branch points in the vasculature as well as features on the leaf margin, instead of just leaf shape alone as in traditional morphometric approaches. Nonetheless, the method overcomes a central problem in the morphometric analysis of leaves: comparing leaves with very different morphologies.

To develop a morphometric method that 1) comprehensively measures shape features in leaves, both locally and globally, 2) can compare disparate leaves shapes, 3) is robust against noise commonly found in leaf shape data (e.g., internal holes because of overlapping leaflets or small defects introduced during imaging and thresholding), and 4) is potentially compatible with other plant phenotyping needs (e.g., measuring the branching architectures of roots and trees, the spatial distributions of plants in ecosystems, or the texture of different pollen types; Mander et al., 2013; 2017; Li et al., 2017b) we used a persistent homology approach. Persistent homology is a topological data analysis method. Topology is the field of mathematics concerned with properties of space preserved under deformations (e.g., bending) but not tearing or re-attaching. Persistent homology measures topological features across the scales of a function (Edelsbrunner and Harer, 2008; Weinberger, 2011; Li et al., 2017b). The compatibility of persistent homology with numerous functions makes it a versatile method that can be tailored for diverse uses (Li et al., 2017a).

Here, we present a morphometric technique based on topology, using a persistent homology framework, to measure the outlines of leaves and classify them by plant family and region in which they were collected. We analyze 182,707 leaves (freely available to download; Chitwood, 2017a), from both published studies and shapes analyzed for the first time, from 141 plant families and 75 sites throughout the world. We first compare the diverse shapes represented in a common morphospace using persistent homology, which captures traditional shape descriptors in a non-linear fashion. Major phylogenetic groups of plants occupy distinct regions of the morphospace and we estimate plant families that have the most and least diverse leaf shapes. Using persistent homology, we then use a linear discriminant analysis to classify leaves by plant family and collection site. Persistent homology predicts both family and collection site at a rate above chance, and predicts leaf family at 2.7 times and collection site at 1.5 times the rate of traditional shape descriptors. Persistent homology is a topological method that can measure and compare diverse leaf shapes from across seed plants and outperforms traditional shape descriptors in classifying plant families and geographic locations.

## Results

### Dataset and a morphospace defined using traditional shape descriptors

To broadly analyze seed plant leaf shape diversity collected from sites throughout the world, we used both published and unpublished data. In total, 182,707 leaves were analyzed (Table 1). Many of these datasets address specific genetic and developmental questions, focusing on genetic variability within a group or closely related species. Leaves were analyzed from the following publications, pre-prints, and authors focusing on specific groups of plants: *Alstroemeria* (2,392 leaves; Chitwood et al., 2012a), apple (9,619 leaves; Migicovsky et al., 2017), *Arabidopsis* (5,101 leaves; AB, RA, CB, ER, BZ), *Brassica* (1,832 leaves; HA, SG, JCP), *Capsicum* (3,277 leaves; TH, AVD), *Coleus* (34,607 leaves; VC, MF, ML), cotton (2,885 leaves; Andres et al., 2017), grapevine and wild relatives (20,121 leaves; Chitwood et al., 2014; 2016a; 2016b; VC, MF, LK, JL, AM), *Hedera* (common ivy, 865 leaves; Martinez et al., 2016), *Passiflora* (3,301 leaves; Chitwood and Otoni, 2017), Poaceae (866 leaves; LC, TG, PK), wild and cultivated potato (1,840 leaves; DF, SJ), tomato and wild relatives (82,034 leaves; Chitwood et al., 2012b; 2012c; 2013), and *Viburnum* (2,422 leaves; Schmerler et al., 2012; MD, EE, SS, ES). We also analyzed two datasets that sample broadly across seed plants and from sites throughout the world. The Leafsnap dataset, with 5,733 leaves, represents mostly tree species of the Northeastern United States, but other groups of plants as well (Kumar et al., 2012). The Climate dataset, with 5,812 leaves total, analyzes the relationship between leaf shape and present climates as indicators of paleoclimate (Huff et al., 2003; Royer et al., 2005; Peppe et al., 2011).

**Table 1:** Leaf counts of datasets.

| Leaf type | Count |
| --- | --- |
| *Alstroemeria* | 2,392 |
| Apple | 9,619 |
| *Arabidopsis* | 5,101 |
| *Brassica* | 1,832 |
| *Capsicum* | 3,277 |
| Climate | 5,812 |
| *Coleus* | 34,607 |
| Cotton | 2,885 |
| Grapevine | 20,121 |
| *Hedera* | 865 |
| Leafsnap | 5,733 |
| *Passiflora* | 3,301 |
| Poaceae | 866 |
| Potato | 1,840 |
| Tomato | 82,034 |
| *Viburnum* | 2,422 |
| Total | 182,707 |

We analyzed all leaves using the traditional shape descriptors circularity, aspect ratio, and solidity (Figure 1). These shape descriptors are simple in the sense that they each measure a very specific aspect of shape, but they are powerful in that they can be applied to any shape, which is not necessarily true of other methods that measure shape more comprehensively (such as landmarks, pseudo-landmarks, and Elliptical Fourier Descriptors). Circularity is a ratio of area to perimeter (true perimeter, excluding holes in the middle of the object) measured as 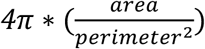 and is sensitive to undulations (like serrations, lobes, and leaflets) along the leaf perimeter, but is also influenced by elongated shapes (like grass leaves) when comparing leaves with such different shapes, as in this analysis. Aspect ratio is measured as (*major axis*)/(*minor axis*) of a fitted ellipse, and it is a robust metric of overall length-to-width ratio of a 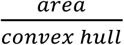 where the convex hull bounds the leaf shape as a leaf. Solidity is measured as polygon. Leaves with a large discrepancy between area and convex hull (such as compound leaves with leaflets, leaves with deep lobes, or leaves with a distinct petiole) can be distinguished from leaves lacking such features using solidity.

**Figure 1:**
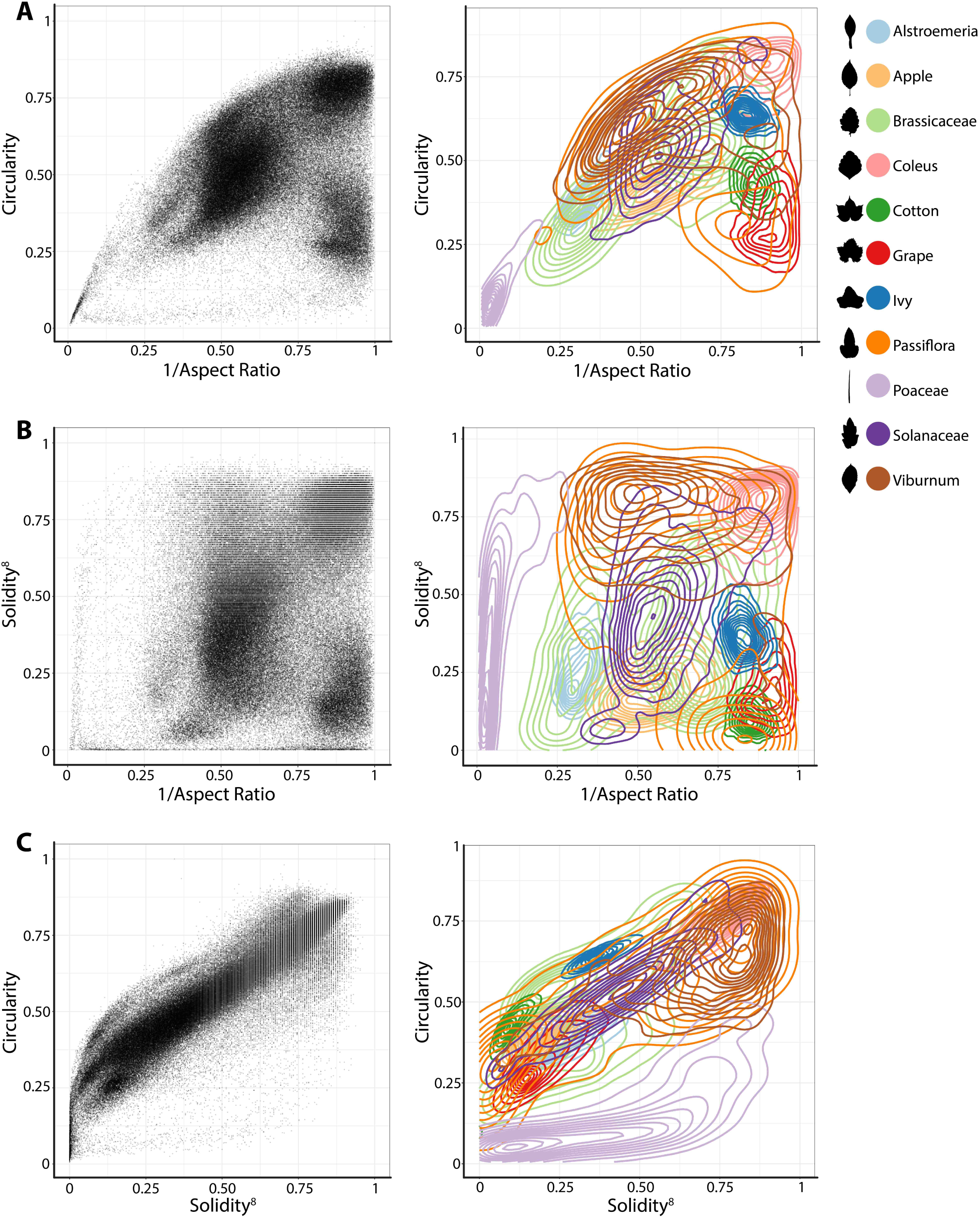
Traditional shape descriptors delimit leaves from different taxonomic groups. **A)** Circularity vs. 1/Aspect Ratio, **B)** Solidity^8^ vs. 1/Aspect Ratio, and **C)** Circularity vs. Solidity^8^. Left: Scatter plots of 182,707 leaves analyzed, from 141 plant families from 75 sites throughout the world. Right: For select taxonomic groups, density plots showing ability of traditional shape descriptors to delimit different leaf shapes and distributions of different groups. Solidity and Aspect Ratio values have been transformed to yield more even distributions. Taxonomic groups are indicated by color and silhouettes of representative leaves close to the overall mean of descriptor values provided.

Differences between groups were visualized as scatterplots and density diagrams (Figure 1), using transformed values of aspect ratio (1/(*aspect ratio*)) and solidity (*solidity*^8^) to create more even distributions that allow the separation between groups to be better visualized. The long leaves of grasses (Poaceae, lavender) are perhaps the most distinct group of leaf shapes. The Brassicaceae (light green) are bimodal in their distribution, reflecting entire vs. highly lobed and compound leaves, as well as differences in petiole length. *Passiflora* (dark orange), Solanaceae (purple), and *Viburnum* (brown) exhibit broad, continuous distributions, which like the Brassicaceae reflect the diversity of leaf shapes in these groups. *Alstroemeria* (light blue), apple (light orange), *Coleus* (pink), cotton (dark green), grapevine (red), and common ivy (dark blue) all have more localized distributions in the morphospace, indicating that shape variation is expressed within a smaller range, relative to other groups, as measured using traditional shape descriptors.

### Persistent homology and non-linear relationships with traditional shape descriptors

Although traditional shape descriptors can describe important shape features across diverse leaves, they do not measure shape comprehensively like landmarks, pseudo-landmarks, and Elliptical Fourier Descriptors. Comprehensive morphometric methods, however, cannot be applied across diverse shapes, only between leaves with similar shapes, as in natural variation studies. We crafted a persistent homology method to quantify the features of leaves, conceptualizing shape as a two-dimensional point cloud of an outline defined by pixels (Li et al., 2017a; Migicovsky et al., 2017). The method begins by calculating a Gaussian density estimator, assigning each pixel a value that indicates the density of neighboring pixels (Figure 2). In leaves, high density pixels with lots of neighbors tend to reside in the sinuses of serrations or lobes or at points of intersection, such as the attachment points of leaflets to the rachis of a compound leaf. Using a Gaussian density estimator, rather than focusing on continuity of a closed contour (as in pseudo-landmarks and Elliptical Fourier Descriptors), minimizes the impact of internal or non-continuous features, such as holes or occlusions made by the overlap of leaflets and lobes (see the bottom palmately-shaped leaf in Figure 2). Sixteen annuli emanating from the centroid of the shape (Figure 2A) serve to partition the leaf into subsets of features, increasing the ability to distinguish between shapes. An annulus kernel for each ring (Figure 2C) is multiplied by the density estimator (Figure 2B) to isolate density features that intersect with the annulus (Figure 2D-E). The resulting density function from each annulus is the function across which topological space is measured. As shown in Figure 2F, beginning with the highest density level, the number of connected features with densities above that level is recorded. Counting the number of connected components minus the number of holes (which is a topological feature, known as the Euler characteristic) continues across the function, proceeding to lower density levels. The value of the curve (y axis in Figure 2F) at each density level (x axis in Figure 2F) records the topological structure across the values of the function, the crux of persistent homology. A curve is recorded for each annulus, so that using our method, the shape of a single leaf is represented by 16 curves.

**Figure 2:**
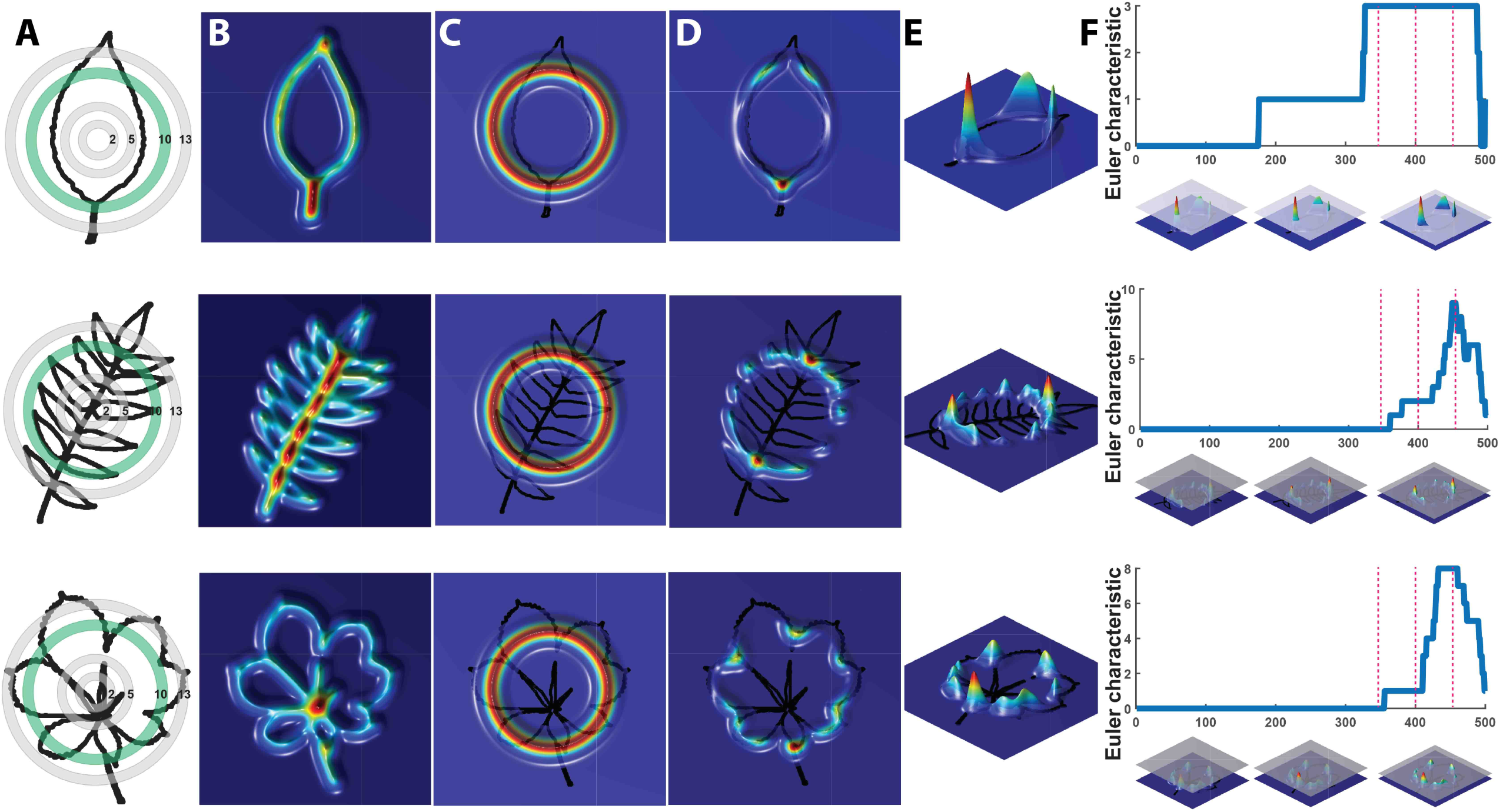
Persistent homology and leaf shape. **A)** Contours of a simple leaf (top), compound pinnate leaf (middle), and compound palmate leaf with a hole and overlap in leaflets (bottom). 16 annuli used to isolate pixel density are shown, with annulus 10 used in subsequent panels indicated in green. **B)** Colormap of a Gaussian density estimator that is robust to noise. Red indicates a larger density of neighboring pixels and blue less density. **C)** An annulus kernel is used to localize and smoothen data. **D)** Multiplication of the annulus kernel with the density estimator isolates density features of the leaf contour. **E)** Side view of the annulus kernel-isolated density features of the leaf. The high peaks in red indicate higher pixel density. **F)** A plane traverses the density function from the highest to lowest densities (x axis). As the plane traverses the function, the topological space is recorded as the number of connected components above the plane at any given point, the Euler characteristic (y axis). Three pink dotted lines correspond to the plane at three points along the density function, which are visualized below the graphs. Together, similar curves from the 16 annuli comprise the persistent homology description of leaf shape.

To analyze the persistent homology output, we discretize each Euler characteristic curve into 500 values (Figure 2F) and then concatenate these values over the 16 annuli, representing each leaf shape as 8,000 values. A Principal Component Analysis (PCA) performed using the 8,000 values creates a leaf morphospace defined by persistent homology (Figure 3). To interpret this morphospace, we colored data using traditional shape descriptor values. Although clear patterns among aspect ratio (Figure 3A), circularity (Figure 3B), and solidity (Figure 3C) with persistent homology data are evident, the relationships are non-linear compared to the orthogonal PC axes. Aspect ratio, circularity, and solidity are similarly correlated with PC1 (rho values of -0.72, 0.70, and 0.61, respectively) demonstrating that persistent homology PCs can capture distinct attributes of shape simultaneously (Figure 3D). The correlations between traditional shape descriptors and persistent homology PCs rapidly diminish among high order PCs (Figure 3D). The non-linear relationship between traditional shape descriptors and persistent homology PCs indicates that persistent homology captures differing combinations of traditional shape descriptors in different ways among the represented leaf shapes. Such non-linear relationships are influenced by the different groups represented in the dataset (Figure 3E). If the Leafsnap and Climate datasets are superimposed as black points on top of a density diagram representing different groups (Figure 3F), then the overall shape of the persistent homology space defined by specific groups is recapitulated. As the Leafsnap and Climate datasets together represent 141 plant families and 75 sites throughout the world, the data suggest that the overall shape and density of the persistent homology morphospace is partially saturated. This does not mean that there is no other significant leaf shape variation to be explored, only that some archetypal leaf shapes are well represented in our dataset. The boundaries of the persistent homology morphospace allow for speculation. Likely the morphospace is 1) bimodal, defined by elongated leaf shapes found in some Poaceae and Gymnosperms (specifically Pinophyta in the Leafsnap and Climate datasets) compared to other leaf shapes and 2) is defined by variation spanning entire to deeply lobed (or even compound) leaf shapes, as represented by *Passiflora*, Solanaceae, and *Vibrunum* across PC1. Of course, other leaf shape variation exists (and is even visually apparent in the plots of PC2 vs. PC1) and other PCs in this dataset remain to be explored. The dataset does not come near to sampling all existing leaf shapes.

**Figure 3:**
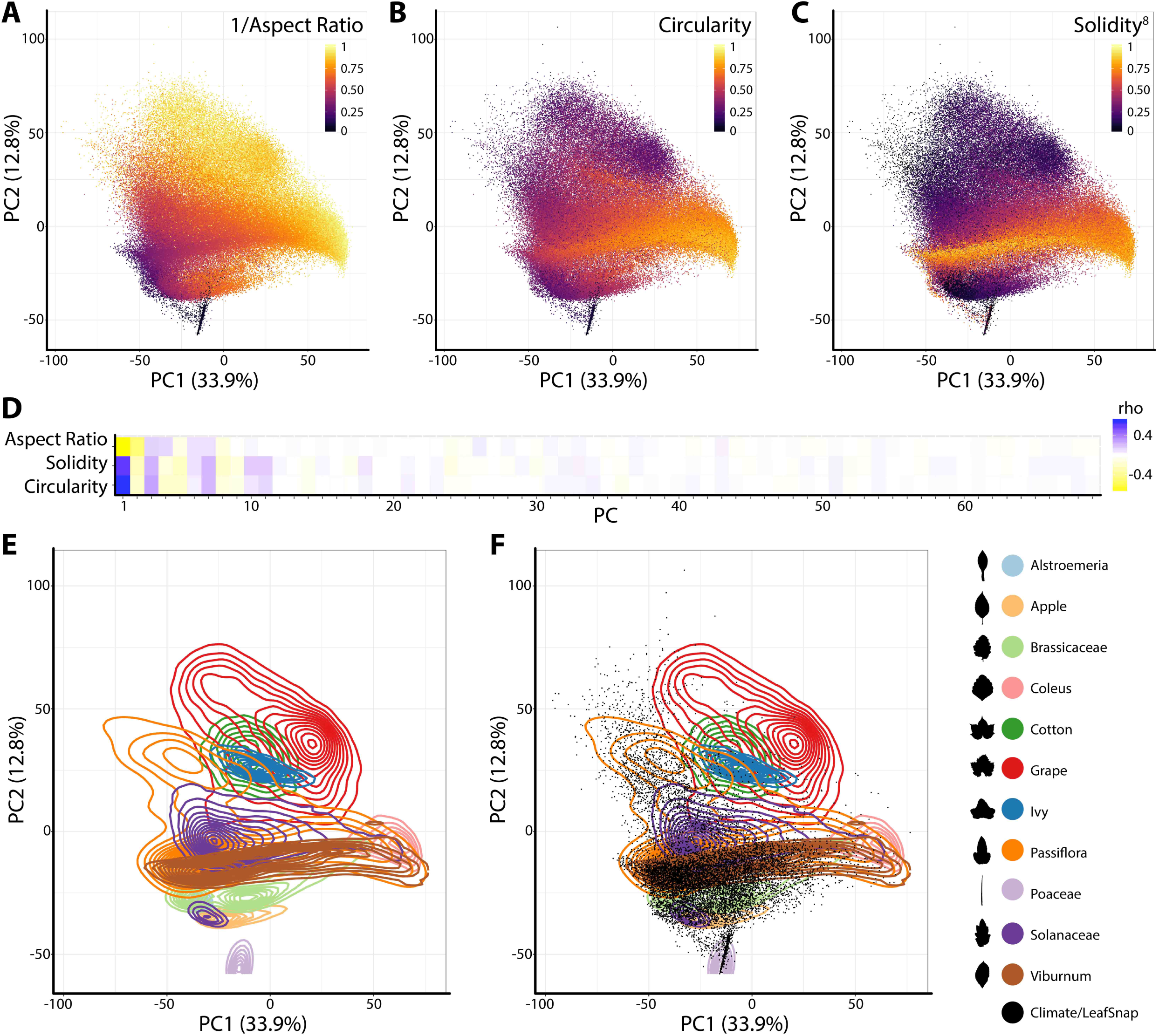
Principal Component Analysis (PCA) of persistent homology results. Principal Component 2 (PC2) vs. PC1 based on persistent homology results for 182,707 leaves colored by **A)** 1/Aspect Ratio, **B)** Circularity, and **C)** Solidity^8^. Aspect Ratio and Solidity values have been transformed to yield more even distributions. Note non-linear relationships between traditional shape descriptors and persistent homology PCs. **D)** Correlations between aspect ratio, circularity, and solidity and PCs 1-69 (representing 90% of variation). Positive and negative Spearman’s rho values are indicated as blue and yellow, respectively. **E)** Density plots show distributions of selected taxonomic groups in persistent homology PCA and **F)** Climate and Leafsnap datasets, representing 141 plant families from 75 sites throughout the world, are superimposed as black dots. Taxonomic groups are indicated by color and silhouettes of representative leaves close to the overall mean of descriptor values provided.

### Differences in leaf shape between phylogenetic groups and the most diverse plant families

We were interested in detecting difference in leaf shape between phylogenetic groups and performed a Principal Component Analysis (PCA) for just the Leafsnap and Climate datasets (Table 1), which together represent 141 plant families, but without the over-representation from specific taxonomic groups presented earlier. Visualizing gymnosperms, magnoliids, rosids I, rosids II, asterids I, and asterids II across PCs 1-10 (representing 73% of shape variance) clear differences in persistent homology shape space can be detected (Figure 4). Differences in shape are most easily detected for the earliest diverging lineages. For example, gymnosperms occupy a distinct region of morphospace defined by PCs 1-6 (Figure 4A-C) compared to angiosperms. Subtler differences between recently diverging groups can also be seen. Asterids II, for example, are excluded from some regions of morphospace occupied by rosids I/II and asterids I for PCs 1-4 (Figure 4A-B).

**Figure 4:**
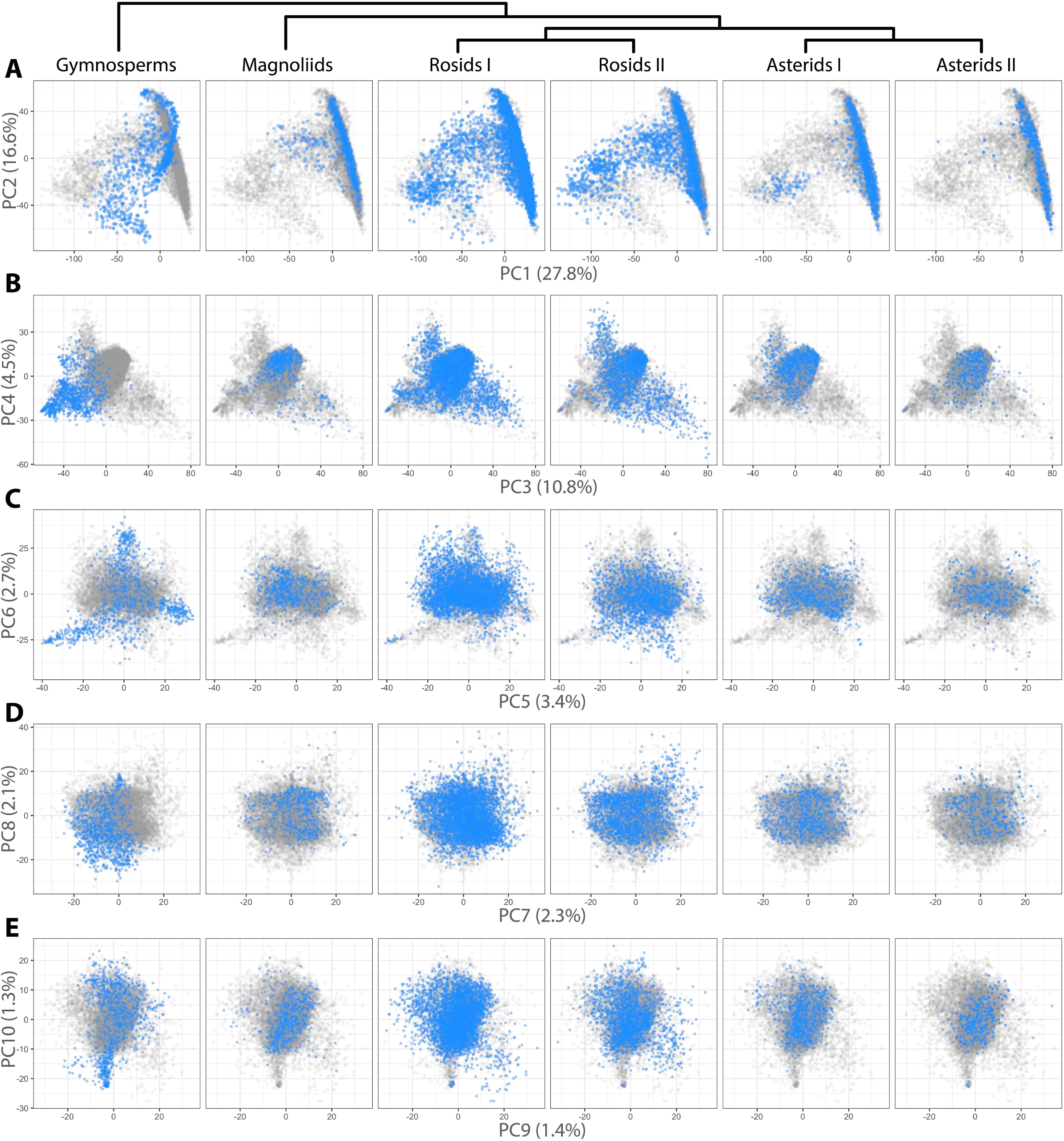
Differences in leaf shape between phylogenetic groups. Gymnosperm, magnoliid, rosid I, rosid II, asterid I, and asterid II leaves (left to right) are each plotted in blue against all samples (gray) for **A)** PC2 vs. PC1, **B)** PC4 vs. PC3, **C)** PC6 vs. PC5, **D)** PC8 vs. PC9, and **E)** PC10 vs. PC9. Percent variance explained by each PC is indicated.

Differences in occupied morphospace between phylogenetic groups prompted us to ask: are plant families diverse across all PCs or just some, and what are the most and least morphologically diverse families? To answer the first question, we calculated variance across PCs 1-179 (representing >95% of all shape variance) for each plant family and then ranked families from most to least variable for each PC (Figure 5A). Visualizing the ranked variability of families across PCs (the most variable ranked families for a PC depicted as yellow, the least variable black, Figure 5A), it is apparent that the most diverse tend be the most diverse across PCs. Increased variability in persistent homology PCs, though, might simply be due to more leaves in some families compared to others. Indeed, the most diverse plant families are also the most represented in our dataset, as seen when families are arranged by abundance (Figures 5A, see bar graph of counts on the right side). Because highly variable families tend to be variable across PCs, we took the median rank of variance across PCs as a measure of overall family leaf shape diversity. The relationship between –median rank variance and log10(count) is linear (Figure S1). Using linear regression, we took the residuals from the model as an estimate of plant family leaf shape diversity, corrected for differences in sample size (Figure 5B). A wilcoxon signed rank test on residuals indicates that asterids I are marginally significant (p = 0.08) for lacking diversity (two sided, mu = 0) but other groups (gymnosperms, p = 0.25; magnoliids, p = 0.20; rosids I, p = 0.97; rosids II, p = 0.63; asterids II, p = 0.63) show no detectable biases in diversity. The overall results indicate that, for the current dataset, leaf shape diversity within major phylogenetic plant groups is equivalent, but specific families have higher estimated leaf shape diversity than others.

**Figure 5:**
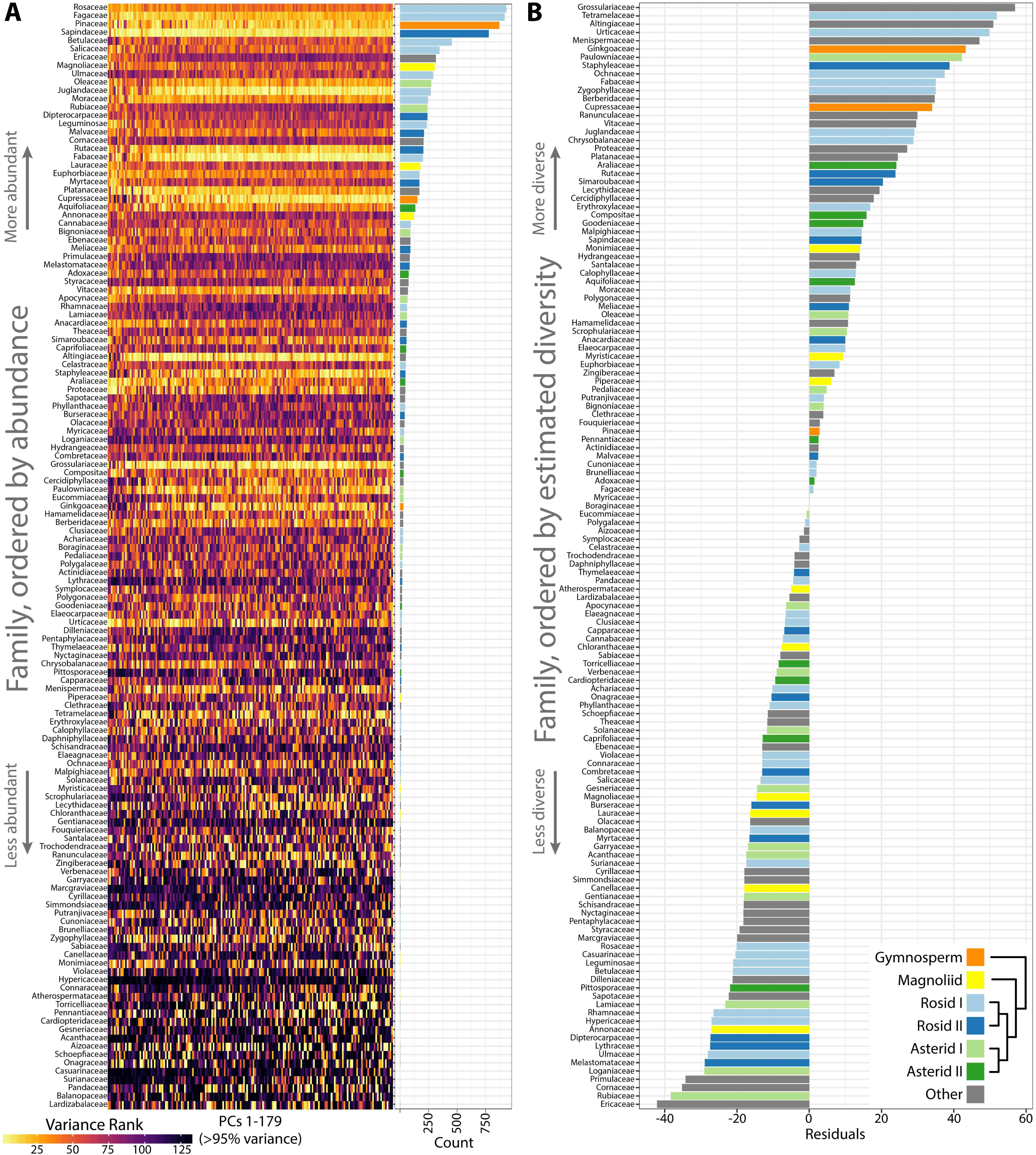
Highly variable plant families are variable across Principal Components (PCs) and estimates of leaf shape diversity by family. **A)** Variance was measured for each plant family and then ranked from most variable (yellow) to least variable (black) for each PC. Plant families are ordered by abundance, as seen in the bar graph (right) indicating count number in the dataset. The most abundant plant families in the dataset tend to be the most variable. **B)** Linear regression was used to model the -median variance ranking for each plant family as a function of log10(count). The residuals are estimates of plant family leaf shape diversity, as corrected for representation in the dataset. Higher residual values indicate higher estimated leaf shape diversity. Gymnosperms, orange; magnoliids, yellow; rosids I, light blue; rosids II, dark blue; asterids I, light green; asterids II, dark green; other groups, gray.

### Persistent homology predicts plant family and region and outperforms traditional shape descriptors

The separation of different groups in the traditional shape descriptor (Figure 1) and persistent homology (Figures 3-4) morphospaces suggests the ability to predict the phylogenetic identity of a leaf based on its shape. Previous computer vision approaches coupled with machine learning have successfully predicted plant family and order using vein patterning and margin features (Wilf et al., 2016). Can the same be done using a persistent homology analysis of the outline alone? Using the Leafsnap and Climate datasets (Table 1) that together represent 141 plant families, we used a Linear Discriminant Analysis (LDA) on PCs 1-179, representing >95% of the persistent homology morphospace variation, to create a classifier scheme. Leaves were then reassigned to the linear discriminant space using a cross-validated “leave one out” approach (Venables and Ripley, 2002) and the results visualized as a confusion matrix (Figure 6), plotting the actual family identity of leaves as a function of the proportion of their predicted family identity. Using persistent homology, there was a 27.3% correct plant family assignment rate of leaves. Using a bootstrapping approach permuting plant family identity against leaf shape information, a 27.3% correct reclassification rate or higher was never achieved in 1,000 bootstrapped simulations, indicating that assignment is above chance. This outperforms traditional shape descriptor prediction (at a rate of 10.2%) by 2.7 times (Table 2), and including both persistent homology and traditional shape descriptor data only marginally increases the prediction rate (to 29.1%) over that of persistent homology alone (27.3%), indicating that persistent homology largely captures the same shape features as traditional descriptors, but provides additional information as well.

**Figure 6:**
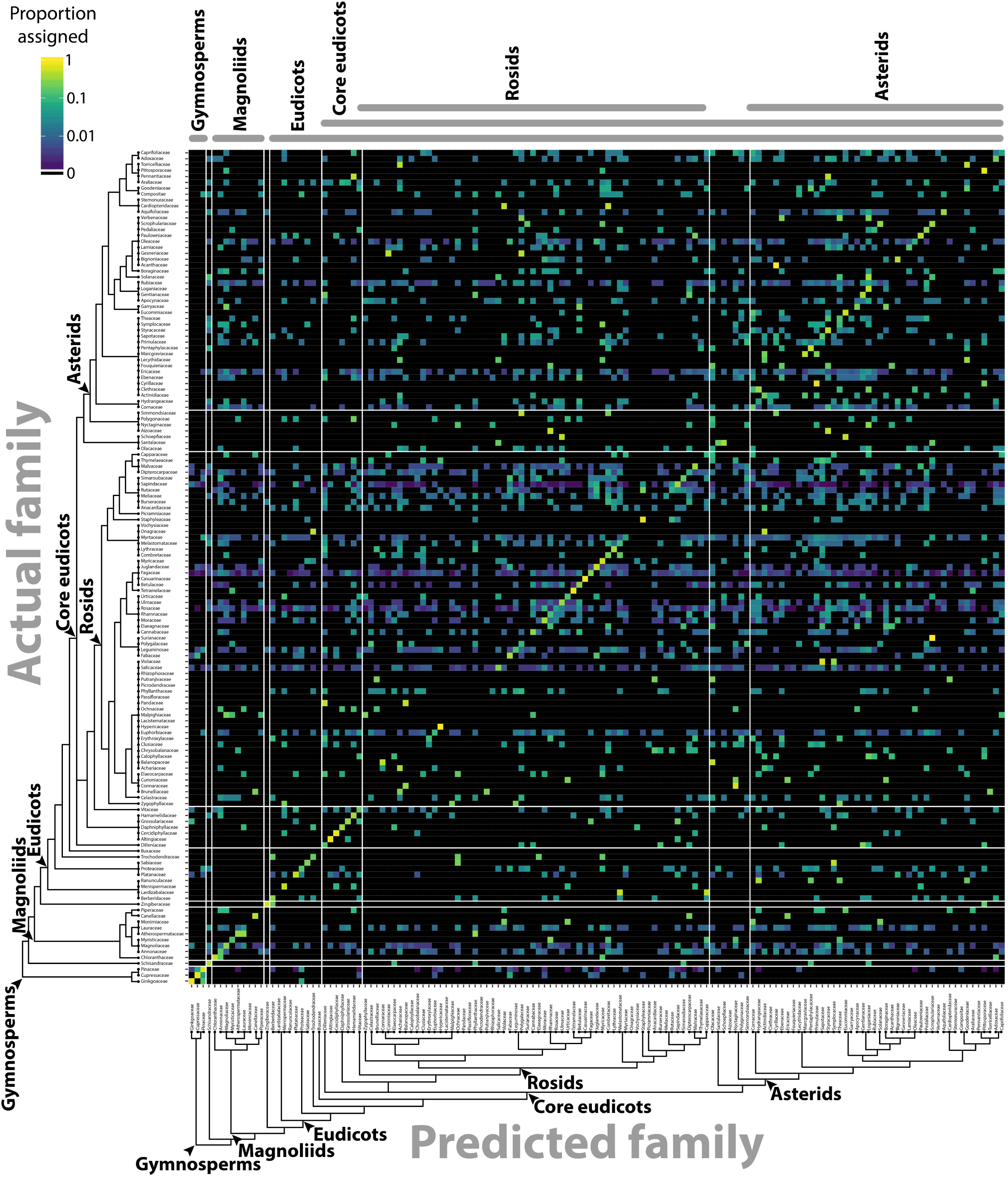
Predicting plant family using persistent homology. Using persistent homology data from the Climate and Leafsnap datasets, a Linear Discriminant Analysis (LDA) was used as a classifier to predict plant family, cross-validated using a jackknifed “leave one out” approach. The vertical axis indicates actual plant family and the horizontal axis predicted plant family. Color indicates proportion of leaves from each actual plant family assigned to each predicted family, such that proportions across the horizontal axis sum to 1. Black indicates no assignment. A phylogeny indicating key taxonomic groups is provided.

**Table 2:** Overall prediction rates of plant family and collection site using different morphometric methods

| Prediction | Datasets | Method | Correct |
| --- | --- | --- | --- |
| Plant family | Climate, Leafsnap | Persistent homology | 27.3% |
| Plant family | Climate, Leafsnap | Traditional descriptors | 10.2% |
| Plant family | Climate, Leafsnap | Both methods | 29.1% |
| Site | Climate | Persistent homology | 14.5% |
| Site | Climate | Traditional descriptors | 9.5% |
| Site | Climate | Both methods | 16.2% |

Previous studies analyzing correlations between leaf shape with present and ancient climates debated the presence of “phylogenetic invariant” features that vary by climate, not phylogenetic context. The Climate dataset includes leaves from 75 sites throughout the world (Table 1). Like the phylogenetic prediction above, we sought to determine the degree that geographic location (regardless of plant family) can be predicted from shape alone. An LDA performed on PCs 1-191, representing >95% of the persistent homology morphospace variation for the Climate dataset, can predict the site where a leaf was collected (Figure 7) at a rate of 14.5% (Table 2). Although much lower than the overall prediction rate by plant family (27.3%), a rate of 14.5% or higher was never achieved in 1,000 bootstrapped simulations, indicating that assignment is above chance. Persistent homology outperforms traditional shape descriptors (at a rate of 9.5%) by 1.5 times (Table 2), and including both persistent homology and traditional shape descriptor data only marginally increases the prediction rate (to 16.2%) over that of persistent homology alone (14.5%).

**Figure 7:**
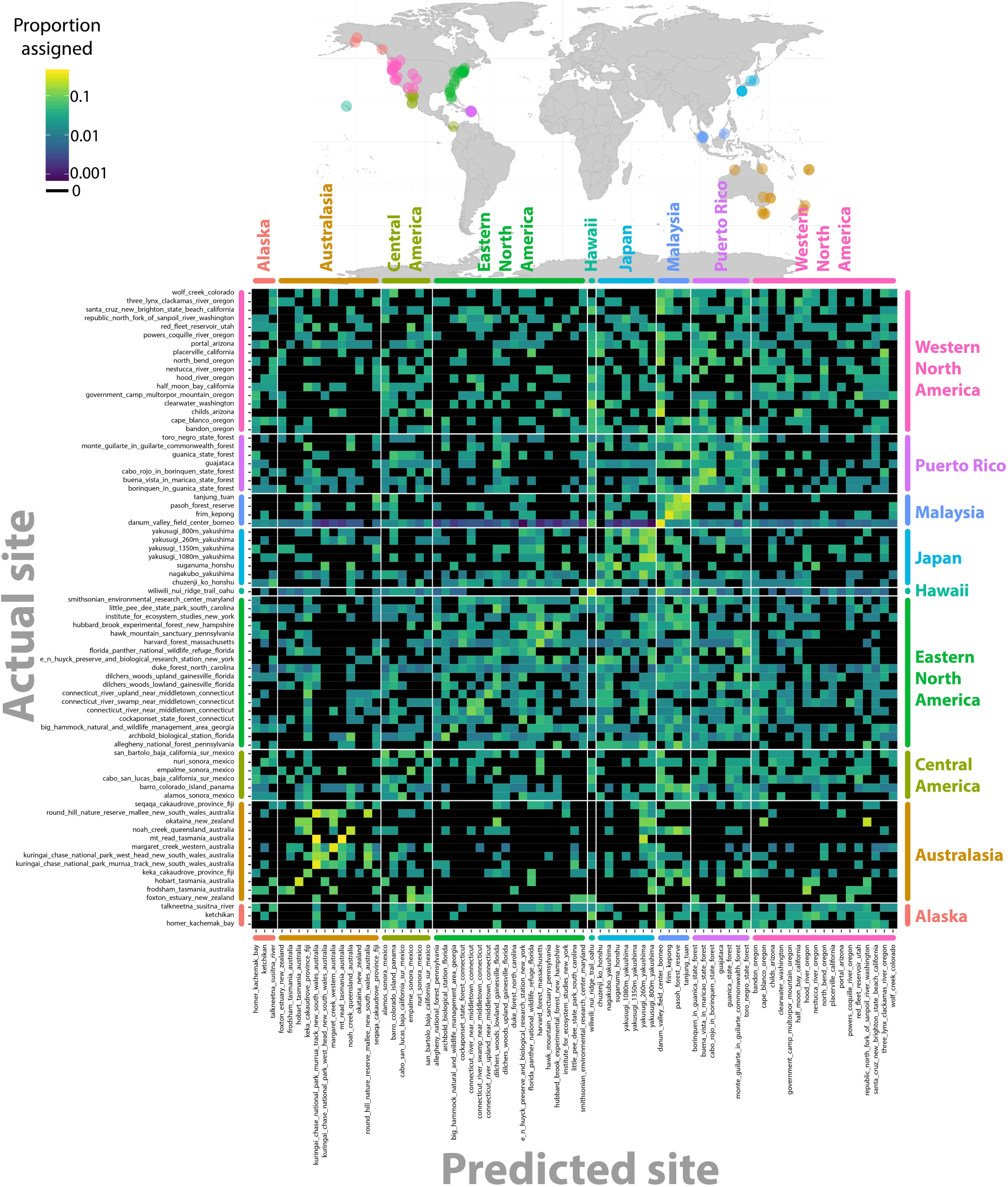
Predicting collection site using persistent homology. Using persistent homology data from the Climate dataset, a Linear Discriminant Analysis (LDA) was used as a classifier to predict collection site, cross-validated using a jackknifed “leave one out” approach. The vertical axis indicates actual collection site and the horizontal axis predicted collection site. Color indicates proportion of leaves from each actual collection site assigned to each predicted collection site, such that proportions across the horizontal axis sum to 1. Black indicates no assignment. Sites are grouped into nine different regions that are indicated by color on a map.

Although the overall prediction rates of 27.3% for plant family and 14.5% for site collected are relatively low (Table 2), it is important to remember that they are above the level of chance (determined by bootstrapping, 1,000 simulations) and that the rates are not evenly distributed across factor levels. Plant family prediction rates vary from 0-100%, and site collected prediction rates vary from 0-40% (Figure 8). The variability in rates is not overly influenced by sampling depth or variation within a group. For example, prediction rate of plant family and abundance are correlated at rho = 0.37, and the correlation with median rank PC variance is rho = -0.24. Although comprehensive, our dataset does not begin to encompass the total shape variation present in a plant family or region and there are undoubtedly collection biases in the data influencing prediction. Other factors than diversity within a group or the degree to which it is sampled, though, likely influence prediction rate too.

**Figure 8:**
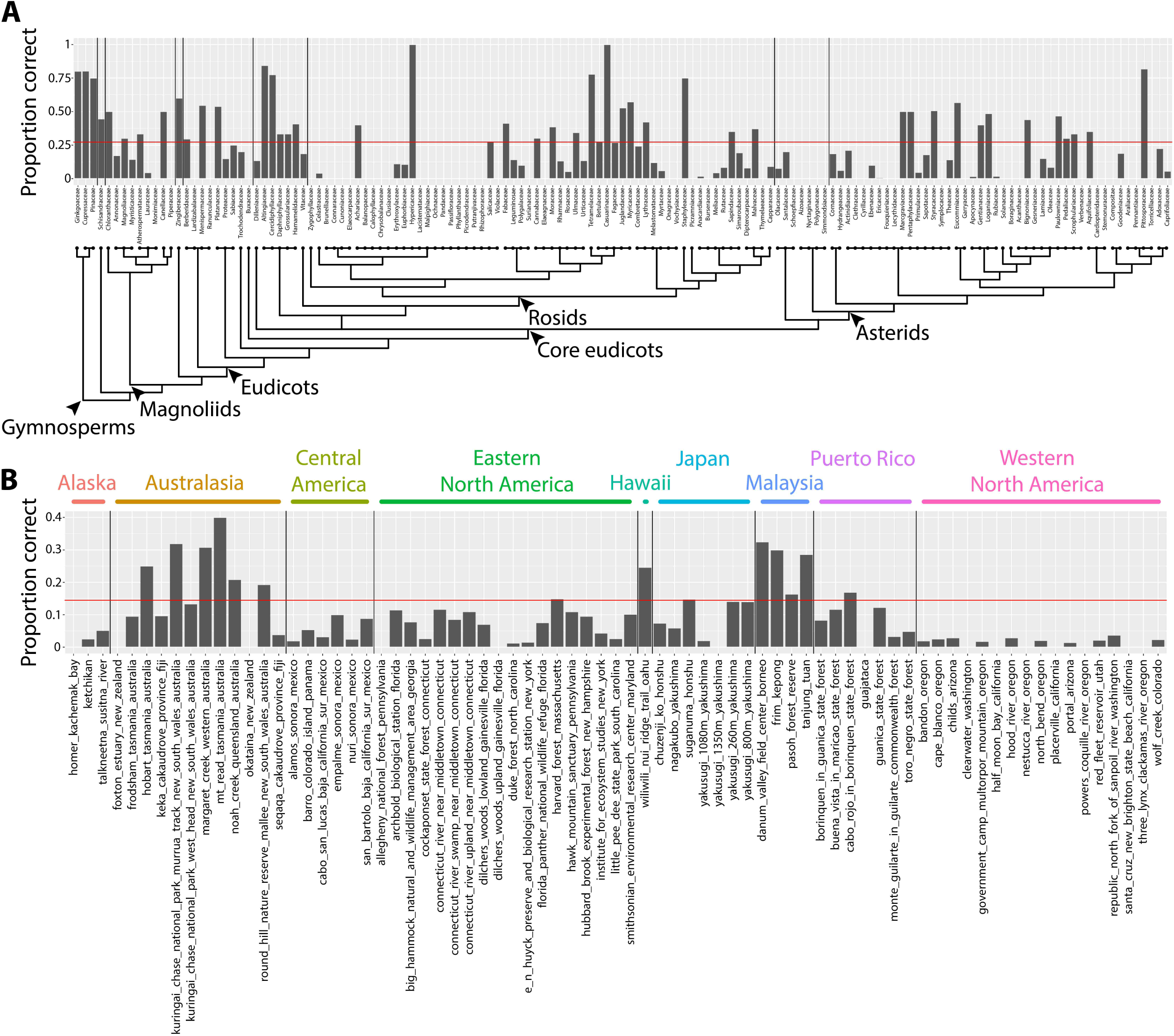
Prediction rates using persistent homology data across plant families and collection sites. **A)** Proportion of leaves from each family correctly assigned. Red line indicates overall correct prediction rate of plant family of 27.3%. Phylogeny and major taxonomic groups are indicated. **B)** Proportion of leaves from each collection site correctly assigned. Red line indicates overall correct prediction rate of collection site of 14.5%. Collection sites are grouped by region, indicated by color.

## Discussion

We have presented a new morphometric method using persistent homology, a topological approach, that can comprehensively measure leaf shape. Other methods measure leaf shape comprehensively, including traditional landmarks, pseudo-landmarks, and Elliptical Fourier Descriptors (EFDs). However, no method comparatively analyzes the diverse *shapes* of leaves in seed plants (simple leaves, deeply lobed leaves, compound leaves of different shapes, leaves with differing numbers of leaflets or lobes, or large variation in petiole length and shape), only naturally varying leaves among related plant species. Other morphometric methods that only analyze the external contour of shapes are sensitive to artifacts, such as internal holes made by the overlap of leaflets or lobes, or small errors during thresholding and isolation. Finally, although appropriate for plant organs that can be represented by discrete shapes—like leaves, petals, seeds, or other lateral organs—current morphometric techniques fail to capture other attributes of plant architecture, like the branching patterns of roots, shoots, and inflorescences. A framework that can not only measure shape, but other features that are important to the plant form, is currently lacking.

By converting shapes into a topological space, as defined by a function that isolates subsets of the shape and describes it in terms of neighboring pixel density (Figure 2), the described persistent homology approach can compare disparate leaf shapes across seed plants, allowing for the approximation of the overall leaf morphospace (Figure 3). By estimating pixel density, the method accommodates internal features (such as holes caused by leaflet overlap) or small processing artifacts, that do not unduly influence the output compared to the absence of such imperfections. The ability to compare shapes broadly and be robust against processing artifacts will enable large scale data analyses in the future, such as the analysis of digitized herbarium vouchers, ecological studies, or genetic and developmental insights into complex morphologies, for which current morphometric approaches are not designed. We detected clear differences in leaf shape between major phylogenetic groups (Figure 4) and estimated leaf shape diversity across plant families (Figure 5), demonstrating that a persistent homology approach is relevant for large-scale morphometric studies across plant evolution. The ability to comprehensively measure shapes permits alternative statistical approaches, moving beyond descriptive statistics used with traditional shape descriptors (Figure 1) and allowing for classifier and prediction approaches (Figures 6-8; Table 2). Theoretically, a unifying morphometric framework that can accommodate not only shapes but the branching architectures of plants, is lacking. As we have previously described, persistent homology functions are ideal to apply to branching plant structures as topological spaces (Li et al., 2017b). The morphometric approach described here applied to leaf shapes is compatible with similar persistent homology methods, creating a shared framework in which the plant form can be measured (Li et al., 2017a).

## Materials and Methods

### Leaf shapes

The 182,707 leaf outlines from 141 plant families from 75 sites throughout the world used in this manuscript are available to download (Chitwood, 2017a). This file directory includes x,y coordinates that form the outlines of the leaves. Separate folders contain text files with x,y coordinates for the leaves from each of the indicated groups in Table 1. Within each folder, original x,y coordinates and scaled coordinates are provided. This dataset contains leaves from both published and unpublished sources (see text for details; Andres et al., 2017; Chitwood et al., 2012a; 2012b; 2012c; 2013; 2014; 2016a; 2016b; Chitwood and Otoni, 2017; Huff et al., 2003; Kumar et al., 2012; Li et al., 2017a; Martinez et al., 2016; Migicovsky et al., 2017; Peppe et al., 2011; Royer et al., 2005; Schmerler et al., 2012; *Arabidopsis* BA, RA, CB, ER, BZ; *Brassica* HA, SG, JCP; *Capsicum* TH, AVD; *Coleus* VC, MF, ML; grapevine and wild relatives VC, MF, LK, JL, AM; Poaceae LC, TG, PK; wild and cultivated potato DF, SJ; *Viburnum* MD, EE, SS, ES).

### Persistent homology

The MATLAB code necessary to recapitulate the persistent homology analysis in this manuscript can be found in the following GitHub repository (Li, 2017): https://github.com/maoli0923/Persistent-Homology-All-Leaf

Persistent homology is a flexible method to quantify branching structures (Edelsbrunner and Harer, 2008; Weinberger, 2011; Li et al., 2017b), point clouds (Ghrist, 2008), two-dimensional and three-dimensional shapes (Gamble and Heo, 2010), and textures (Mander et al., 2013; 2017). Each of these different phenotypes can be described by a multidimensional vector (e.g. Euler characteristic curve), integrating how homology (e.g. path-connected components) persists across the scales of a tailored mathematical function.

Leaf contours are represented as two-dimensional point clouds extracted from binary images (Figure 2A). We use a Gaussian density estimator, which can be directly derived from the point cloud and is also robust to noise, to estimate the neighborhood density of each pixel. Denser point regions, such as serrations, lobes, or the attachment points of leaflets, have higher function values (Figure 2B). Formally, the Gaussian density estimator is defined as 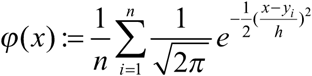, where 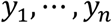 are the data points and *h* is a bandwidth parameter. Because a set of local and regional topologies may often be more effective to represent shapes, we use a local persistent homology technique to subset the density estimator into 16 concentric annuli centered around the centroid of the leaf (Figures 2A, D). To achieve this, we multiply this function by a “bump” function K which highlights an annulus, defined as 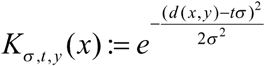, where *y* is the center of the annulus, *tσ* determines its radius, and the parameter *σ* is its width (Figure 2C). Each local function emphasizes the density function falling in the annulus. Given a threshold and a local function, the points whose function values are greater than this threshold form a subset (superlevel set). Changing this threshold value from the maximum function value to its minimum value, we can get an expanding sequence of subsets, or a superlevel set filtration. Figure 2E shows the shapes above a plane, an example of a superlevel set filtration. For each subset, we calculate the Euler characteristic, which equals the number of connected components minus the number of holes. Thus, for a sequence of subsets, we get a sequence of numbers (a multidimensional vector). All 16 annuli derive 16 multidimensional vectors which are concatenated into an overall vector used for analysis. Principal Component Analysis (PCA) was performed in MATLAB on the vectors and PC scores and percent variance explained by each PC used in subsequent analyses.

### Statistical analysis and visualization

The R code (R Core Team, 2017) and data necessary to recapitulate the statistical analyses and figures in this manuscript can be found as a zipped folder directory on figshare (Chitwood, 2017b): https://figshare.com/articles/LeafMorphospace/4985561/1

Unless otherwise specified, all graphs were visualized using ggplot2 (Wickham, 2016). Scatterplots were visualized using the geom_point() function, density plots were visualized with the geom_density2d() function, heatmaps were visualized using the geom_tile() function, and colors were selected from ColorBrewer (Harrower and Brewer, 2003) and viridis (Garnier, 2017). Other visualization functions used and specific parameters that can be found in the code used to generate the figures (Chitwood, 2017b).

Variance was calculated for each plant family for each principal component using var() and families ranked for each principal component using rank() (Figure 5). Linear regression was performed using lm() and residuals retrieved to estimate leaf shape diversity for each plant family (Figure S1). The Wilcoxon signed rank test was performed using wilcox.test() to test for higher or lower than expected phylogenetic diversity using a two-sided test with mu = 0. Linear Discriminant Analysis (LDA) was performed using the lda() function in the package MASS (Venables and Ripley, 2002). LDA was performed using the number of principal components that contributed at least 95% of all variance in each analysis (PCs 1-179 for phylogenetic prediction and PCs 1-191 for site prediction). The Leafsnap and Climate datasets were used for phylogenetic prediction (Figure 6) whereas just the Climate dataset was used for site prediction (Figure 7). Prediction using the discriminant space was performed using CV = TRUE for a “leave one out” cross-validated jack-knifed approach and the priors set equal across factor levels. Both the phylogenetic and site LDA prediction rates were bootstrapped over 1,000 simulations. A for loop was used, permuting family or site identity against leaf identity, performing an LDA on the permuted data, and recording the correct prediction rate for each permuted simulation. For both the phylogenetic and site predictions, a permuted correct prediction rate (out of 1,000 simulations) higher than the actual correct prediction rate was never detected.

## Supplemental figure legend

**Supplemental Figure 1:**
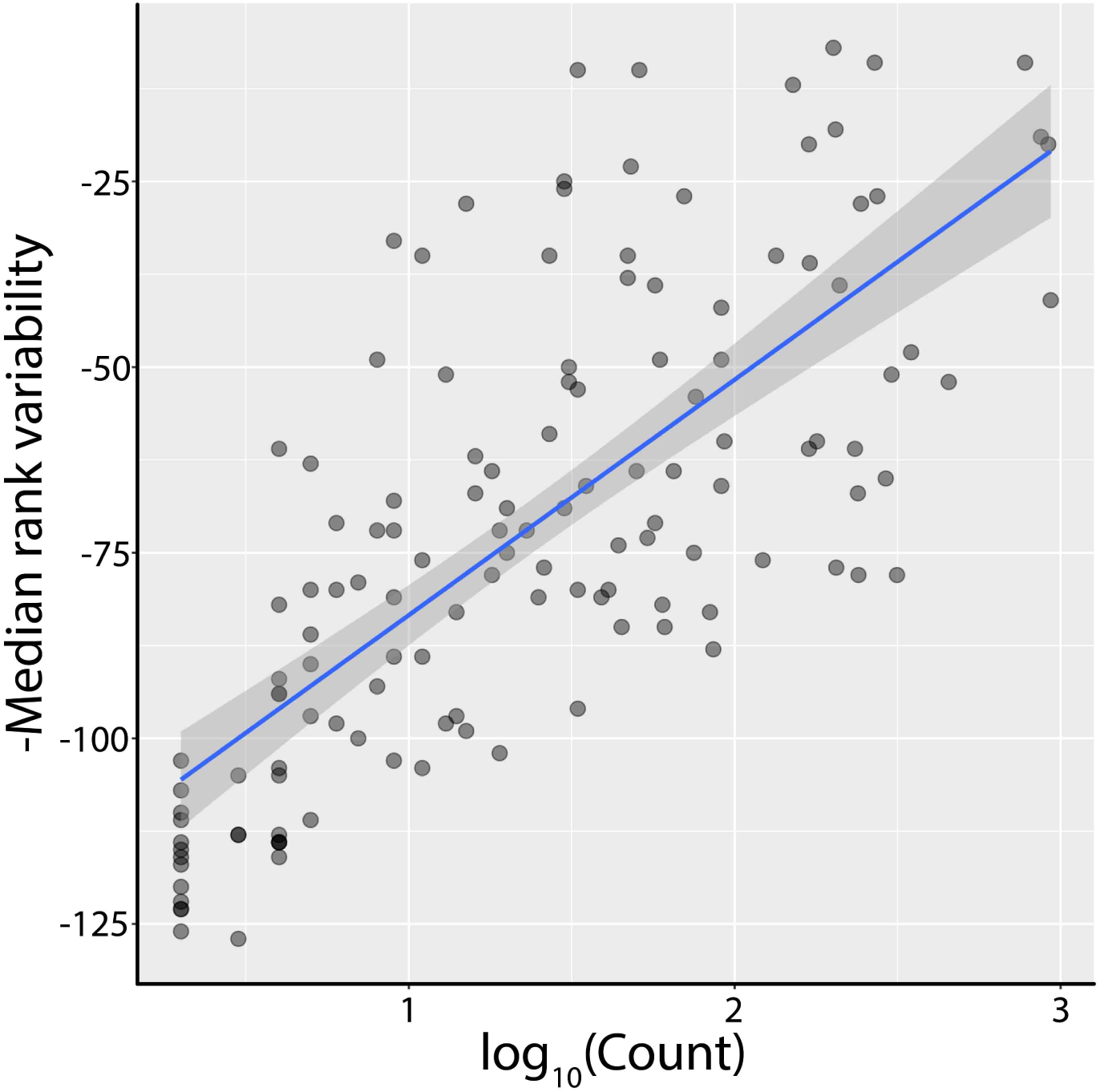
Linear relationship between median ranked variability and count. Linear regression was used to model -median rank variability (higher values indicated more variability within a plant family) as of function of the abundance of each plant family in the dataset, as measured by log_10_(leaf count). The model (shown in blue) was used to estimate overall leaf shape variance in plant family, as corrected for sampling depth, by using the residuals from the model as an indication of diversity.

## References

Andres RJ, Coneva V, Frank MH, Tuttle JR, Samayoa LF, Han SW, Kaur B, Zhu LL, Fang H, Bowman DT, Rojas-Pierce M. Modifications to a *LATE MERISTEM IDENTITY1* gene are responsible for the major leaf shapes of Upland cotton (*Gossypium hirsutum* L.). Proceedings of the National Academy of Sciences of the United States of America. 2017 Jan 3;114(1):E57-66.

Bensmihen S, Hanna AI, Langlade NB, Micol JL, Bangham A, Coen ES. Mutational spaces for leaf shape and size. HFSP Journal. 2008 Apr 1;2(2):110-20.

Bookstein FL. Morphometric tools for landmark data: geometry and biology. Cambridge University Press; 1997 Jun 28.

Chitwood DH, Naylor DT, Thammapichai P, Weeger AC, Headland LR, Sinha NR. Conflict between intrinsic leaf asymmetry and phyllotaxis in the resupinate leaves of *Alstroemeria psittacina*. Frontiers in Plant Science. 2012a Aug 10;3:182.

Chitwood DH, Headland LR, Filiault DL, Kumar R, Jiménez-Gómez JM, Schrager AV, Park DS, Peng J, Sinha NR, Maloof JN. Native environment modulates leaf size and response to simulated foliar shade across wild tomato species. PLoS One. 2012b Jan 12;7(1):e29570.

Chitwood DH, Headland LR, Kumar R, Peng J, Maloof JN, Sinha NR. The developmental trajectory of leaflet morphology in wild tomato species. Plant Physiology. 2012c Mar 1;158(3):1230-40.

Chitwood DH, Kumar R, Headland LR, Ranjan A, Covington MF, Ichihashi Y, Fulop D, Jiménez-Gómez JM, Peng J, Maloof JN, Sinha NR. A quantitative genetic basis for leaf morphology in a set of precisely defined tomato introgression lines. The Plant Cell. 2013 Jul 1;25(7):2465-81.

Chitwood DH, Ranjan A, Martinez CC, Headland LR, Thiem T, Kumar R, Covington MF, Hatcher T, Naylor DT, Zimmerman S, Downs N. A modern ampelography: a genetic basis for leaf shape and venation patterning in grape. Plant Physiology. 2014;164(1):259-72.

Chitwood DH, Klein LL, O'Hanlon R, Chacko S, Greg M, Kitchen C, Miller AJ, Londo JP. Latent developmental and evolutionary shapes embedded within the grapevine leaf. New Phytologist. 2016a Apr 1;210(1):343-55.

Chitwood DH, Rundell SM, Li DY, Woodford QL, Tommy TY, Lopez JR, Greenblatt D, Kang J, Londo JP. Climate and developmental plasticity: interannual variability in grapevine leaf morphology. Plant Physiology. 2016b Mar 1;170(3):1480-91.

Chitwood DH, Otoni WC. Morphometric analysis of Passiflora leaves: the relationship between landmarks of the vasculature and elliptical Fourier descriptors of the blade. Gigascience. 2017 Jan 1;6(1):1-13.

Chitwood DH. Leaf_coordinates. Figshare. 2017a. Accessed June 17, 2017. https://doi.org/10.6084/m9.figshare.5056441.v1

Chitwood DH. LeafMorphospace. Figshare. 2017b. Accessed May 29, 2017. https://doi.org/10.6084/m9.figshare.4985561.v1

Edelsbrunner H, Harer J. Persistent homology-a survey. Contemporary mathematics. 2008 Feb 29;453:257-82.

Freeman H. Computer processing of line-drawing images. ACM Computing Surveys (CSUR). 1974 Mar 1;6(1):57-97.

Gamble J, Heo G. Exploring uses of persistent homology for statistical analysis of landmark-based shape data. Journal of Multivariate Analysis. 2010 Oct 31;101(9):2184-99.

Garnier S. viridis: Default Color Maps from ‘matplotlib’. R package version 0.4.0. 2017 https://CRAN.R-project.org/package=viridis

Ghrist R. Barcodes: the persistent topology of data. Bulletin of the American Mathematical Society. 2008;45(1):61-75.

Gower JC. Generalized procrustes analysis. Psychometrika. 1975 Mar 27;40(1):33-51.

Harrower M, Brewer CA. ColorBrewer. org: an online tool for selecting colour schemes for maps. The Cartographic Journal. 2003 Jun 1;40(1):27-37.

Huff PM, Wilf P, Azumah EJ. Digital future for paleoclimate estimation from fossil leaves? Preliminary results. Palaios. 2003 Jun;18(3):266-74.

Kuhl FP, Giardina CR. Elliptic Fourier features of a closed contour. Computer Graphics and Image Processing. 1982 Mar 1;18(3):236-58.

Kumar N, Belhumeur P, Biswas A, Jacobs D, Kress WJ, Lopez I, Soares J. Leafsnap: A computer vision system for automatic plant species identification. Computer Vision–ECCV 2012. 2012:502-16.

Langlade NB, Feng X, Dransfield T, Copsey L, Hanna AI, Thébaud C, Bangham A, Hudson A, Coen E. Evolution through genetically controlled allometry space. Proceedings of the National Academy of Sciences of the United States of America. 2005 Jul 19;102(29):10221-6.

Li M, Frank MH, Coneva V, Mio W, Topp CN, Chitwood DH. Persistent homology: a tool to universally measure plant morphologies across organs and scales. bioRxiv. 2017a https://doi.org/10.1101/104141

Li M, Duncan K, Topp CN, Chitwood DH. Persistent homology and the branching topologies of plants. American Journal of Botany. 2017b 104(3):349-353.

Li M. Persistent-Homology-All-Leaf. GitHub. 2017. Accessed May 29, 2017. https://github.com/maoli0923/Persistent-Homology-All-Leaf

Mander L, Li M, Mio W, Fowlkes CC, Punyasena SW. Classification of grass pollen through the quantitative analysis of surface ornamentation and texture. Proceedings of the Royal Society of London B: Biological Sciences. 2013 Nov 7;280(1770):20131905.

Mander L, Dekker SC, Li M, Mio W, Punyasena SW, Lenton TM. A morphometric analysis of vegetation patterns in dryland ecosystems. Royal Society Open Science. 2017 Feb 1;4(2):160443.

Martinez CC, Chitwood DH, Smith RS, Sinha NR. Left–right leaf asymmetry in decussate and distichous phyllotactic systems. Philosophical Transactions of the Royal Society 2016 Dec 19;371(1710):20150412.

Migicovsky Z, Li M, Chitwood DH, Myles S. Morphometrics reveals complex and heritable apple leaf shapes. bioRxiv. 2017 https://doi.org/10.1101/139303

Peppe DJ, Royer DL, Cariglino B, Oliver SY, Newman S, Leight E, Enikolopov G, Fernandez-Burgos M, Herrera F, Adams JM, Correa E. Sensitivity of leaf size and shape to climate: global patterns and paleoclimatic applications. New Phytologist. 2011 May 1;190(3):724-39.

R Core Team. R: A language and environment for statistical computing. R Foundation for Statistical Computing. 2017. Vienna, Austria. Accessed May 29, 2017. https://www.R-project.org/

Royer DL, Wilf P, Janesko DA, Kowalski EA, Dilcher DL. Correlations of climate and plant ecology to leaf size and shape: potential proxies for the fossil record. American Journal of Botany. 2005 Jul 1;92(7):1141-51.

Schmerler SB, Clement WL, Beaulieu JM, Chatelet DS, Sack L, Donoghue MJ, Edwards EJ. Evolution of leaf form correlates with tropical–temperate transitions in Viburnum (Adoxaceae). Proceedings of the Royal Society of London B: Biological Sciences. 2012 Oct 7;279(1744):3905-13.

Venables WN, Ripley BD. Modern Applied Statistics with S. Springer; 2017 New York

Weinberger S. What is… persistent homology? Notices of the AMS. 2011 Jan;58(1):36-9.

Wickham H. ggplot2: elegant graphics for data analysis. Springer; 2016 Jun 8 New York

Wilf P, Zhang S, Chikkerur S, Little SA, Wing SL, Serre T. Computer vision cracks the leaf code. Proceedings of the National Academy of Sciences of the United States of America. 2016 Mar 7:201524473.

